# Otolith microchemistry identifies diadromous populations of Patagonian river fishes

**DOI:** 10.1101/174656

**Authors:** Dominique Alò, Cristián Correa, Horacio Samaniego, Corey A. Krabbenhoft, Thomas F. Turner

## Abstract

**Compliance with Ethical Standards:** Otolith analysis was funded by a RAC grant from the University of New Mexico, USA. The Government of Chile supported the drafting of this document with a CONICYT Doctoral Fellowship to D. Alò in 2015 and to C. Correa through grants CONICYT-PAI N°82130009, and FONDECYT-Iniciación en la Investigación N°11150990.

All applicable international, national, and/or institutional guidelines for the care and use of animals were followed. Specimens were collected under permits No. 3587, 29 December 2006, and No. 2886, 4 November 2008 (amendment No. 602, 12 February 2009) granted by the Chilean Subsecretary of Fishing to C. Correa. The McGill University Animal Care Committee (UACC), Animal Use Protocol No. 5291, approved use and handling of animals.

**Abstract:** Coastal habitats in Chile are hypothesized to support a number of diadromous fishes. The objective of this study was to document migratory life histories of native galaxiids and introduced salmonids from a wide latitudinal range in Chilean Patagonia (39-48°S). Otolith microchemistry data were analysed using a recursive partitioning approach to test for diadromy. Based on annular analysis of Sr:Ca ratios, a diadromous life history was detected for populations of native *Aplochiton taeniatus, A. marinus*, and *Galaxias maculatus*. Lifetime residency in freshwater was suggested for populations of *A. zebra* and *G. platei*. Among introduced salmonids, populations of *Oncorhynchus tshawytscha* and *O. kisutch* exhibited anadromous migratory patterns, whereas the population of *O. mykiss* screened appeared restricted to freshwater. *Salmo trutta* exhibited variable habitat use consistent with establishment of an ocean-type life history in some populations. The capacity and geographic scope of hydropower development is increasing and may disrupt migratory routes of diadromous fishes. Identification of diadromous species is a critical first step for preventing their loss due to hydropower development.

## Introduction

Only 47 native and 27 non-native inland fish species are currently recognized in Chile, and about 30% of these are thought to exhibit some tolerance for shifting between saline and freshwater habitats (Dyer, 2000; Habit & Victoriano, 2005; Habit et al., 2006; Vila et al., 2011; Ministerio del Medio Ambiente, 2013; Vargas et al., 2015). Furthermore, roughly 15% of these fishes are hypothesized to display diadromous migratory behaviour (Table S1), compared to less than 1% for fishes worldwide (Nelson, 2006).

The term diadromy describes regular, predictable, and physiologically mediated movements between freshwater and the sea. Diadromy necessitates profound physiological changes (i.e., osmoregulation) when shifting from marine to freshwater habitats and vice versa (Gross et al., 1988). Diadromy is obligatory for many populations within a species, but also can be facultative (Dingle & Drake, 2007). The direction of migration depends on life history stages and habitats where reproductive and feeding events occur. The combination of these factors defines three different types of diadromy: anadromy, catadromy, and amphidromy (Myers, 1949; Gross, 1987; Gross et al., 1988; McDowall, 1992, 1997; Limburg et al., 2001) (in particular refer to McDowall 1997 for a review of the terminology and a visual aid).

Given the high percentage of fishes in Chile hypothesized to exhibit some form of diadromy, migration might play an important, yet unrecognized role in establishing national priorities of aquatic biodiversity conservation. At present, a high percentage of the continental ichthyofauna in Chile is categorized with some degree of conservation threat by Chilean environmental agencies and other authors, although conservation categories can be incongruent and threats underestimated (Habit & Victoriano, 2005; Diario Oficial de la Republica de Chile, 2008; Ministerio del Medio Ambiente, 2013; IUCN, 2015; Vargas et al., 2015).

Coastal habitats in Chile appear well suited to support establishment of diadromous species. Andean rivers that flow into the Pacific Ocean include a variety of different habitats in a limited longitudinal distance (average 145 km), spanning from areas of rocky substrates, high gradient, clear waters and low temperatures, to areas of low flow, sandy substrates, and aquatic vegetation (Habit & Victoriano, 2005; Instituto Nacional de Estadisticas, 2015). Spatial habitat heterogeneity is essential for maintenance and completion of diadromous life cycles, and for maintaining evolutionary potential (i.e., genetic diversity) for life history variation (Pulido, 2007; Dingle, 2014). Therefore, fragmentation events imposed by human-made barriers may affect fish fitness and restrict movement between habitats more so than in other areas (Waples et al., 2007).

Patagonian fishes offer a unique opportunity to understand migration patterns in relatively pristine habitats, and establish a baseline against which future potential impacts associated with river impoundments can be compared. Despite strong economic growth and efforts to develop hydroelectric potential to meet the country’s high-energy requirements (Joo et al., 2015), many rivers in southern Chile are still free-flowing, offering opportunities to study pre-impoundment patterns of diadromous migration. For example, galaxiid fishes are distributed across the Southern Hemisphere and diadromy seems to be a recurrent trait among many of the species (McDowall, 1971, 1988). Likewise, salmonids are among some of the best-studied diadromous fishes in the Northern Hemisphere and are now well-established in southern Chile (McDowall, 2002; Correa & Gross, 2008).

Using micro-geochemical data obtained from otoliths, this study sought to determine whether native galaxiids and introduced salmonids exhibit diadromy in Chilean rivers. Otoliths are calcified deposits in the inner ear of fishes that accumulate in ring-like fashion over ontogenetic growth. Elemental analysis of otoliths can help to distinguish origins of marine and freshwater fishes among locations with variable water chemistry. Differing chemical composition of the otolith from the primordium (core) to the edge indicates the different environments in which a fish has lived and allows for hypothesis tests related to patterns of fish movement. When analysed sequentially across an otolith sagittal section, changes in elemental ratios can indicate fine-scale patterns of movement, connectivity, dispersal, and the location of natal habitats (Halden et al., 2000; Howland et al., 2001; Kraus, 2004; Ashford et al., 2005; Campana, 2005; Arkhipkin et al., 2009). To quantify these changes in Patagonian fishes, we applied univariate recursive partitioning approaches based on Classification and Regression Trees (CART) to detect discontinuities in ratios that indicate habitat shifts (Vignon, 2015).

## Methods

### Fish Collections

Between 2004 and 2011, specimens of *Aplochiton zebra, A. taeniatus, A. marinus, Galaxias maculatus, G. platei, Oncorynchus tshawytscha, O. kisutch, O. mykiss,* and *Salmo trutta* were collected using various methods from 6 locations across a large latitudinal range (39.5–48.0° S) in western Patagonia, Chile (Fig. 1, Table 1). At each location, fish specimens were euthanized by an overdose of anaesthetic solution (tricaine-methanesulfonate MS-222 or clove oil). Due to the difficulties in morphological identification, genetic data were used to identify individuals in the genus *Aplochiton* to the species level (Alò et al., 2013).

**Table 1:**
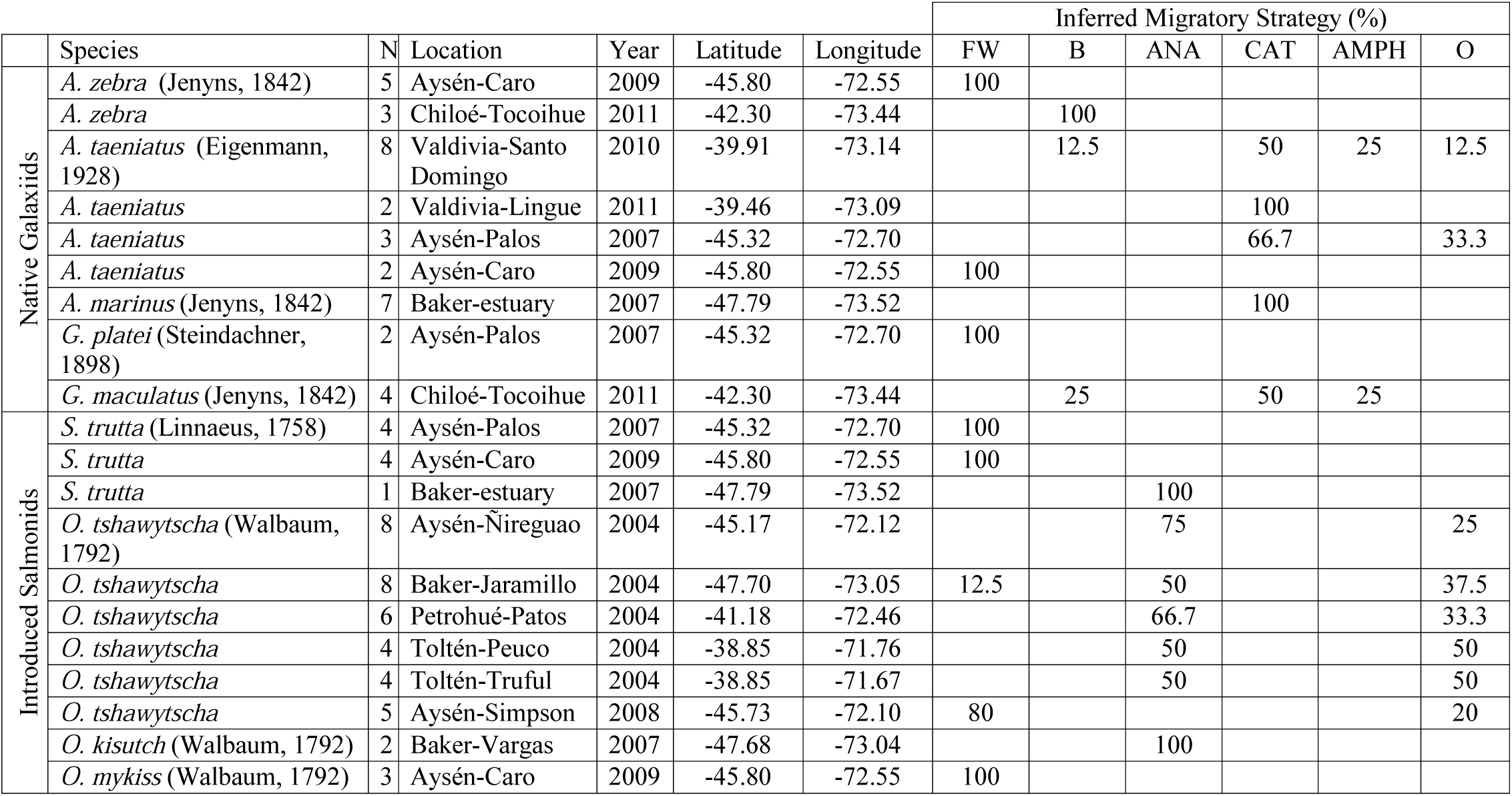
Description of the samples studied and summary of results of migration pattern determination. Values correspond to the percentage of individuals assigned to one of five possible patterns: freshwater resident (FW), brackish water resident (B), anadromous (ANA), catadromous (CAT), amphidromous (AMPH) or else omitted from interpretation due to uncertainties in the otolith transects (O). For individual results, see Table S2

**Fig. 1:**
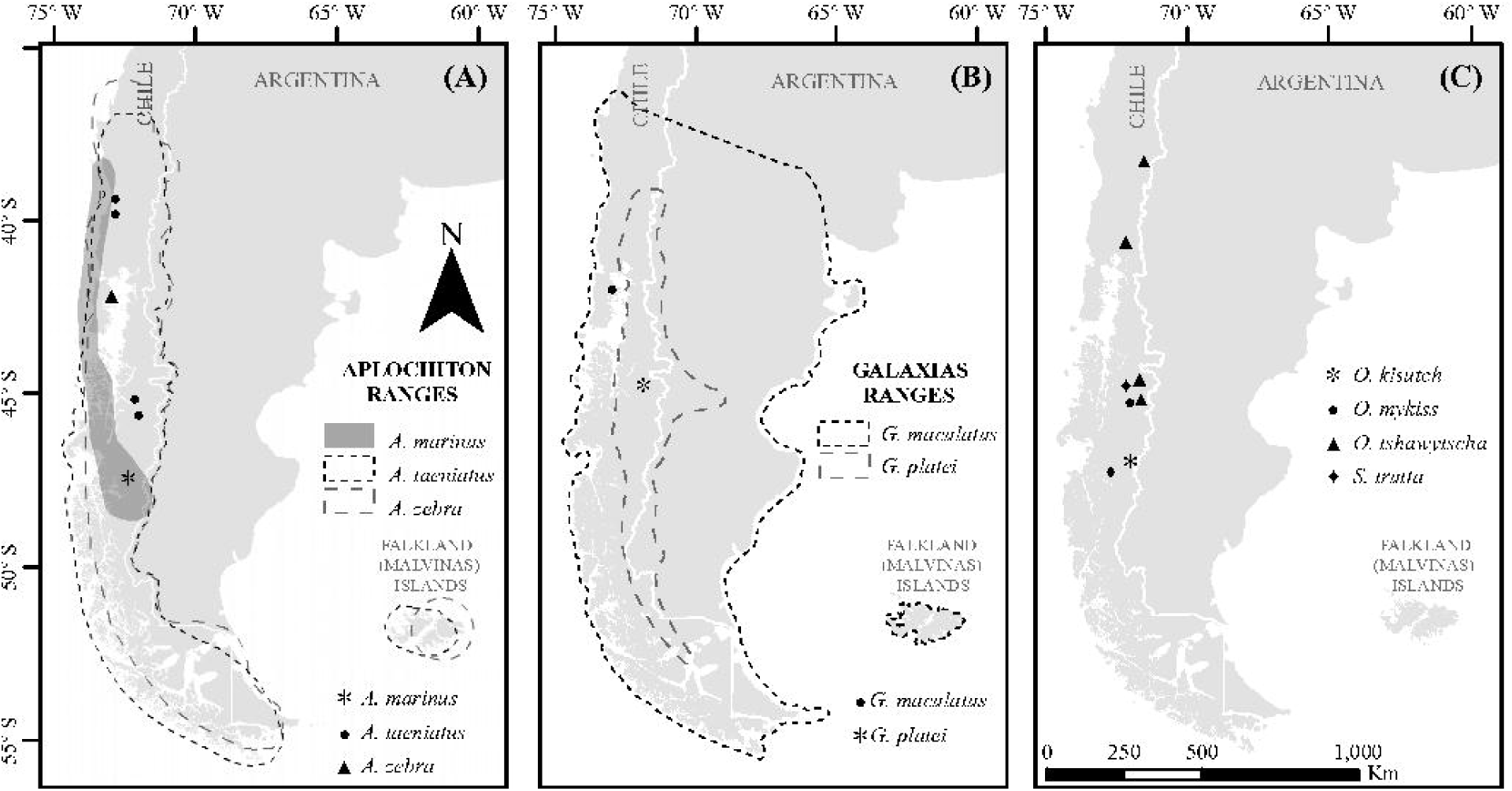
Estimated distribution range for native Chilean galaxiids (shaded and dash lined polygons) and the sampling locations of specimens used in this study (dots) for a) genus *Aplochiton*, b) genus *Galaxias*, c) non-native fishes examined in this study.

### Otolith preparation

Prior to specimen preservation, sagittal otoliths were extracted and either stored dry in test tubes or in 95% ethanol, as elemental compositions and structures of otoliths are not strongly affected by ethanol for the elements assayed (Proctor & Thresher, 1998).

In the laboratory, otoliths were polished, cleaned, and mounted individually on clean glass slides using a thermoplastic cement (Crystalbond™). In order to expose growth rings, 3M™ (fine) and Nanolap ^®^ Technologies (coarse) diamond lapping film wetted with deionized water was used to polish otoliths by hand until a satisfactory sagittal section of annuli was visible. For *Aplochiton* spp., *Galaxias* spp., *S. trutta, O. kisutch* and *O. mykiss* otoliths, a 30-µm and then 3-µm lapping film was used to expose annuli and get a finished polish. *O. tshawytscha* otoliths required larger lapping film (45 and 60 µm) to reach an appropriate view, but were finished with 3 µm film for increased clarity. After the initial polish, and where necessary (largely for *O. tshawytscha*), the adhesive was melted and the otolith flipped for double polishing to produce a thinner section.

Following polishing, the mounting adhesive was dissolved in 100% acetone bath and sonicated for 10 minutes. Larger otoliths were cleaned a second time with acetone as needed. Each otolith was then sonicated twice in Milli-Q water for 5 to 10 minutes each. Following cleaning, otoliths were rinsed a final time in Milli-Q water, transferred to clean vials and placed in a positive laminar flow hood for 24-48 hours to dry.

Acid-washed porcelain forceps were used to mount clean, dry otoliths on acid-washed microscope slides. Otoliths were grouped according to diameter and mounted 10-28 per slide accordingly. Each otolith was placed within one small drop of fresh Crystalbond melted onto a single slide.

Slides were securely kept in acid-washed, sealed petri dishes for transport to Woods Hole Oceanographic Institute (Woods Hole, MA, U.S.A.). There, laser ablation was conducted from October 8^th^ to 11^th^, 2012 (*Aplochiton* spp., *S. trutta, Galaxias* spp., *O. kisutch,* and *O. mykiss*) and again from February 4^th^ to 5^th^, 2013 (*O. tshawytscha*). Laser ablation was performed with a large format laser ablation cell on a New Wave UP193 (Electro Scientific Industries, Portland, Oregon) short pulse width excimer laser ablation system. This was coupled with a Thermo Finnigan Element2 sector field argon plasma spectrometer (Thermo Electron Corporation, Bremen, Germany) for elemental analysis. The laser was configured for single pass, straight line scanning at a speed of 5 µm per second. The laser beam spot size was 50 µm at 75% intensity and 10 Hz pulse rate. Concentrations were determined for elemental Strontium (Sr) and Calcium (Ca) since strontium has been identified as a useful trace element to reconstruct environmental history of fishes and discern habitat shifts across salinity gradients (Secor & Rooker, 2000; Campana, 2005; Pracheil et al., 2014).

Certified standards FEBS-1 and NIES-022 were run before and after each block of 10 to 28 otoliths to account for quality assurance in the measurements (Yoshinaga et al., 2000; Sturgeon et al., 2005). Each otolith was visualized on screen and the intended ablation transect of each sample was plotted digitally and analysed by ablation with a laser beam (refer to Fig. 2 for a visual example). For accuracy of readings, each data point was produced from an average of ten consecutive reads. The ideal double life-history transect ran across each sagittal otolith and through the primordium, thus providing two similar (redundant) patterns related to life history variation, one on either side of the primordium. Interpretations were based on the analyses of both sides of each double transect, if possible. However, in a number of cases, transects were imperfect due to damaged otoliths or inaccurate ablation pathways (Table S2). In such cases, data were analysed as partial transects that can still be used to differentiate diadromous or resident signals (Fig. 2).

**Fig. 2:**
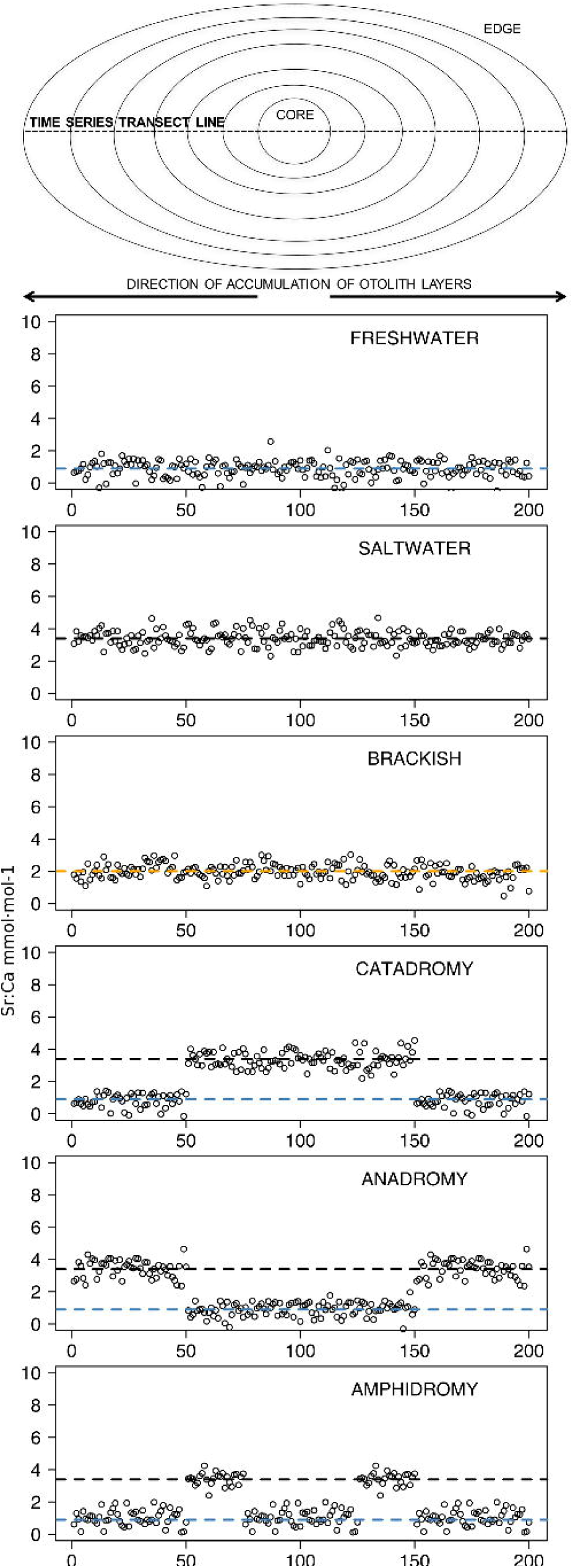
The uppermost image depicts a schematic representation of an otolith, showing how growth rings accrue over time around the core and culminate at the edge. The number of distinct layers in the otolith depends on the age of the individual. The images below represent idealized time series data obtained by repeatedly measuring (via laser ablation and spectrometry) elemental strontium to calcium (Sr:Ca) ratios across the otolith. Each box shows an expected time series for each life-history strategy. The blue dashed line represents the average Sr:Ca for freshwater as 0.9 mmol·mol^−1^, the orange dashed line for estuarine areas at 2.0 mmol·mol^−1^, and the black dashed line is drawn at 3.4 mmol·mol^−1^ as the saltwater mean (Secor & Rooker, 2000)

## Data Analysis

Classification and Regression Trees (CART, Breiman et al. 1984)) were used to detect shifts in elemental ratios across the otolith transect. CART is an alternative to qualitative methods traditionally used to interpret the chronological signal in otolith microchemistry transects (Vignon, 2015). The position along the otolith transect (predictor variable) was recursively partitioned using regression trees in order to differentiate segments of the transect that shared similar mean Sr:Ca values (response variables) (Breiman et al., 1984; Therneau & Atkinson, 1997; De’ath, 2002; Strobl, 2009). CART was implemented in the Tampo library (version 1.0) for R Statistical Software 3.0.2 (Vignon, 2015).

Elemental data and otolith transects were rigorously checked, and outliers caused by recording errors were removed (additive outliers, R Package “tsoutliers” v: 0.6–5, L’opez-de-Lacalle 2016). Summary statistics of Sr:Ca ratios were calculated across all individuals. Since the main goal of this work was to represent movement patterns at a broad scale, CART analysis was used in a semi-supervised manner to identify the presence or absence of sudden discontinuities in the Sr:Ca otolith signal (Vignon, 2015). This was done by introducing three progressively relaxed conditions to the splitting procedure, which required setting a minimum difference in mean values in order to allow a split in elemental signals. The minimal differences required to allow regression tree pruning were set to condition =1.0, cond.=0.7 and cond. =0.5. The detection of one or more discontinuities or splits in the Sr:Ca signal was interpreted as evidence for diadromy, or otherwise, evidence for residency. When diadromy was detected, the direction of ontogenetic movements was inferred from differences in segment means; increasing values indicated movements towards the sea, and vice versa. Further inference about habitat occupancy (freshwaters, estuaries, or the sea) required a visual, heuristic examination of Sr:Ca profiles in relation to published reference values. Finally, all evidence was assembled to make individual inferences about specific migration patterns (amphidromous, catadromous, or anadromous). In this process, otolith transect quality affected our confidence on interpretations; from maximum confidence on inferences from transects that conformed to the model in Fig. 2, to uncertain interpretations from incomplete or faulty transects.

Reference values for the three major habitat types (freshwater, estuarine, and marine) were obtained from a meta-analysis of Sr:Ca profiles from otoliths of 41 fish species in three salinity regimes, and were summarised as follows (extracted from Fig. 7 in Secor and Rooker 2000): Freshwater (salinity 0-5 ppt): 10 species screened, mean Sr:Ca (10-90^th^ percentile range) 0.9 (0.3-1.8) mmol mol^−1^; Estuarine (5-25 ppt): 11 spp., 2.0 (0.9-3.1) mmol mol^−1^; Marine (>25 ppt): 20 spp., 3.4 (1.9-5.2) mmol mol^−1^.

## Results

Observed variation in Sr:Ca ratios was generally bounded within previously reported reference ranges determined by meta-analysis (Fig. 3; Secor & Rooker 2000). CART analysis identified patterns of change in Sr:Ca elemental ratios consistent with the different migratory life histories proposed in the schematic representation in Fig. 2, and some representative individuals for each species are shown in Fig. 4. Details on the splitting results of all individuals under different stringency conditions are given in Fig. 5 whereas Table S2 reports details on the mean and standard deviation at each split. A summary of the inferred migratory strategy for each species is shown in Table 1.

**Fig. 3:**
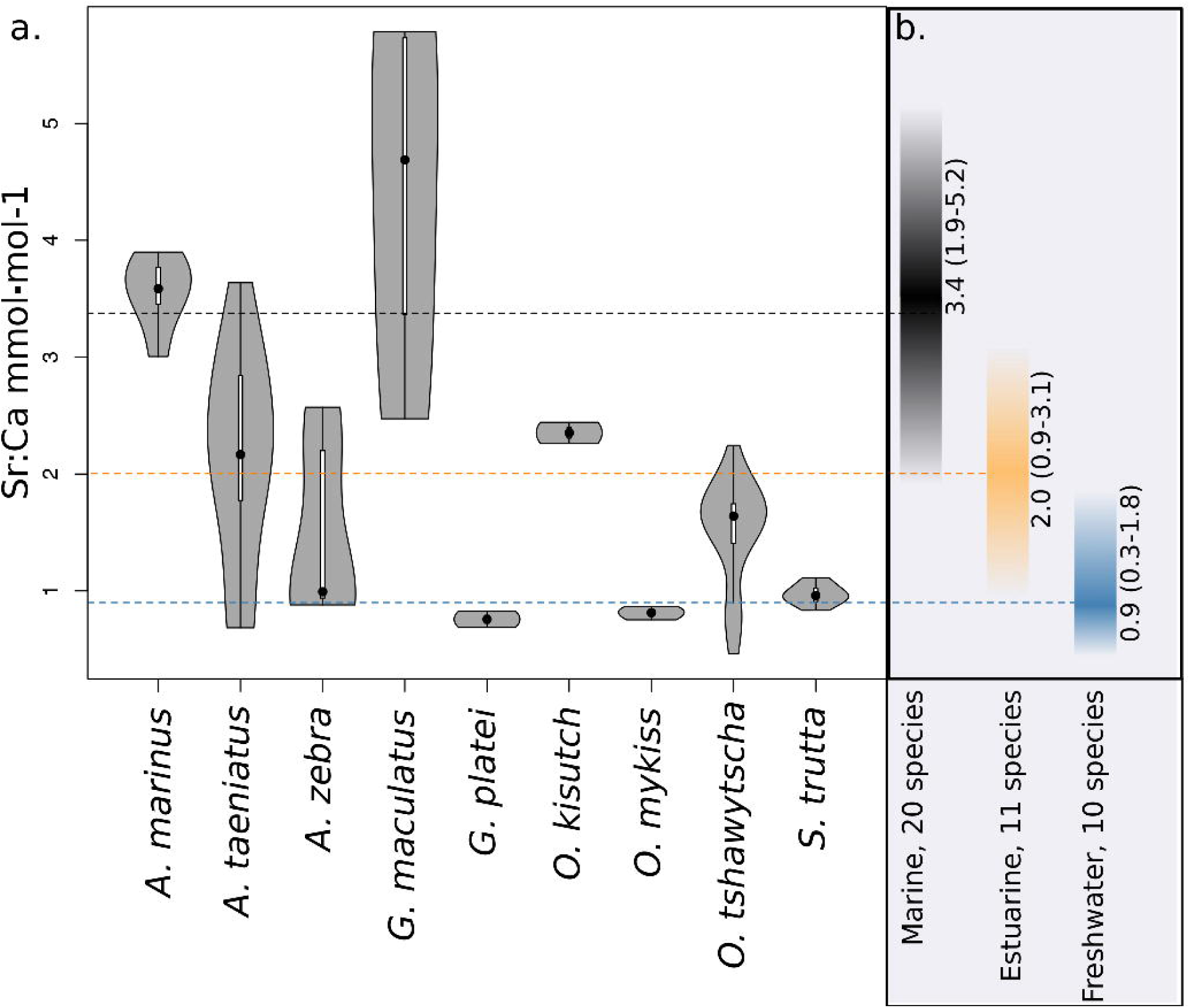
a) Violin plot of Sr:Ca values for otoliths grouped by species. Black dot = median, White line = first to third quartile. Grey areas = kernel density plot. b) Extracted reference values from Fig. 7 in Secor and Rooker 2000. Average values of Sr:Ca ratios for freshwater, brackish and marine water are represented as dashed blue, orange, and black lines, respectively. Fading shades around each mean correspond to 10-90^th^ percentiles for each average value.

**Fig. 4:**
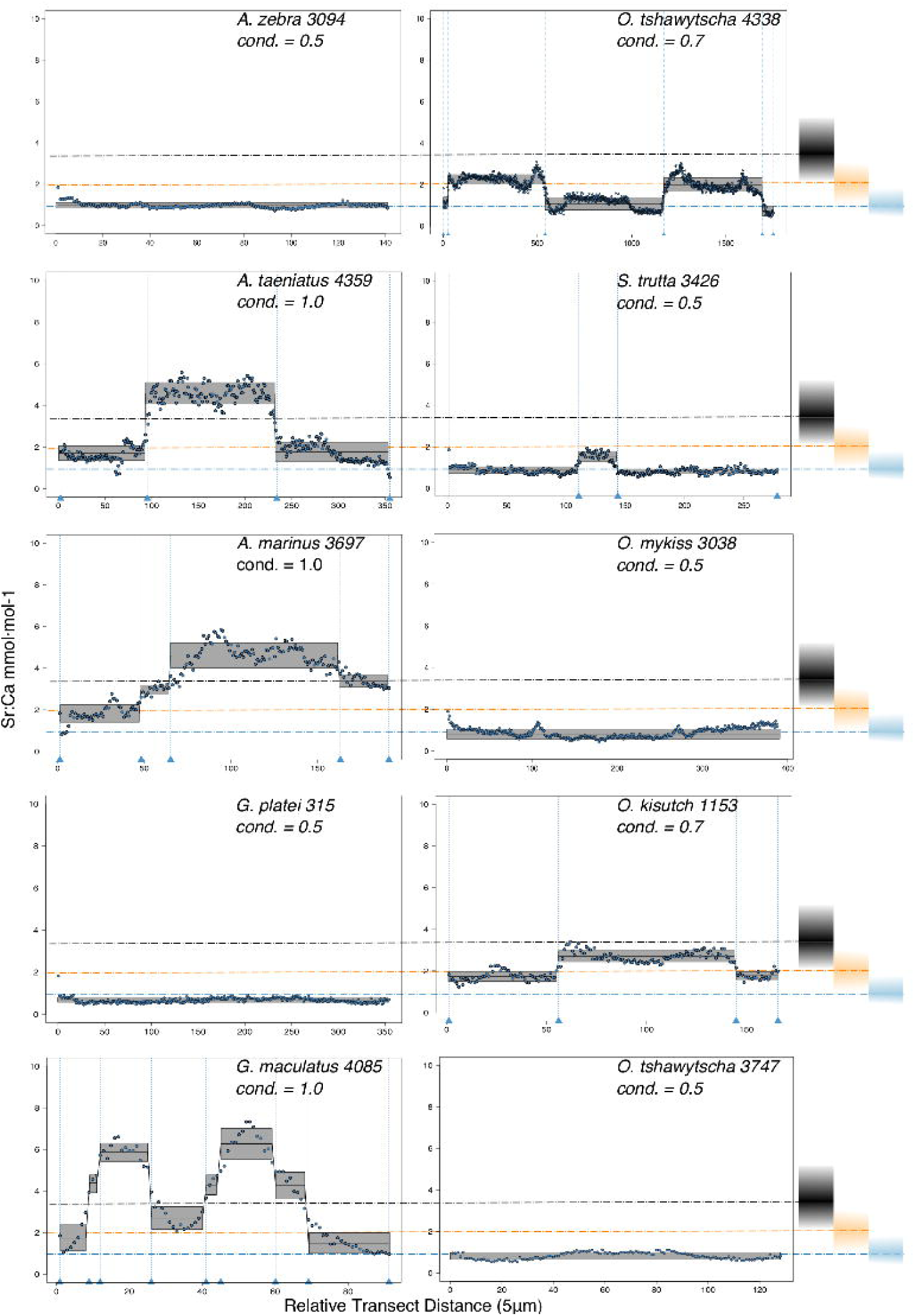
Detection of discontinuities by semi-supervised CART employed on Sr:Ca ratios for representative individuals of native and exotic fishes in southern Chile. Numbers after taxonomic names refer to the individual ID of each fish. The mean and standard deviation are delimited for each cluster in a grey box. Vertical dashed lines indicate splitting points induced by the condition used to fit the regression trees, which is reported for each individual graph as “cond”. Reference values from Secor and Rooker (2000) are reported on the right-hand side of the graph; see legend of Fig. 3.

**Fig. 5:**
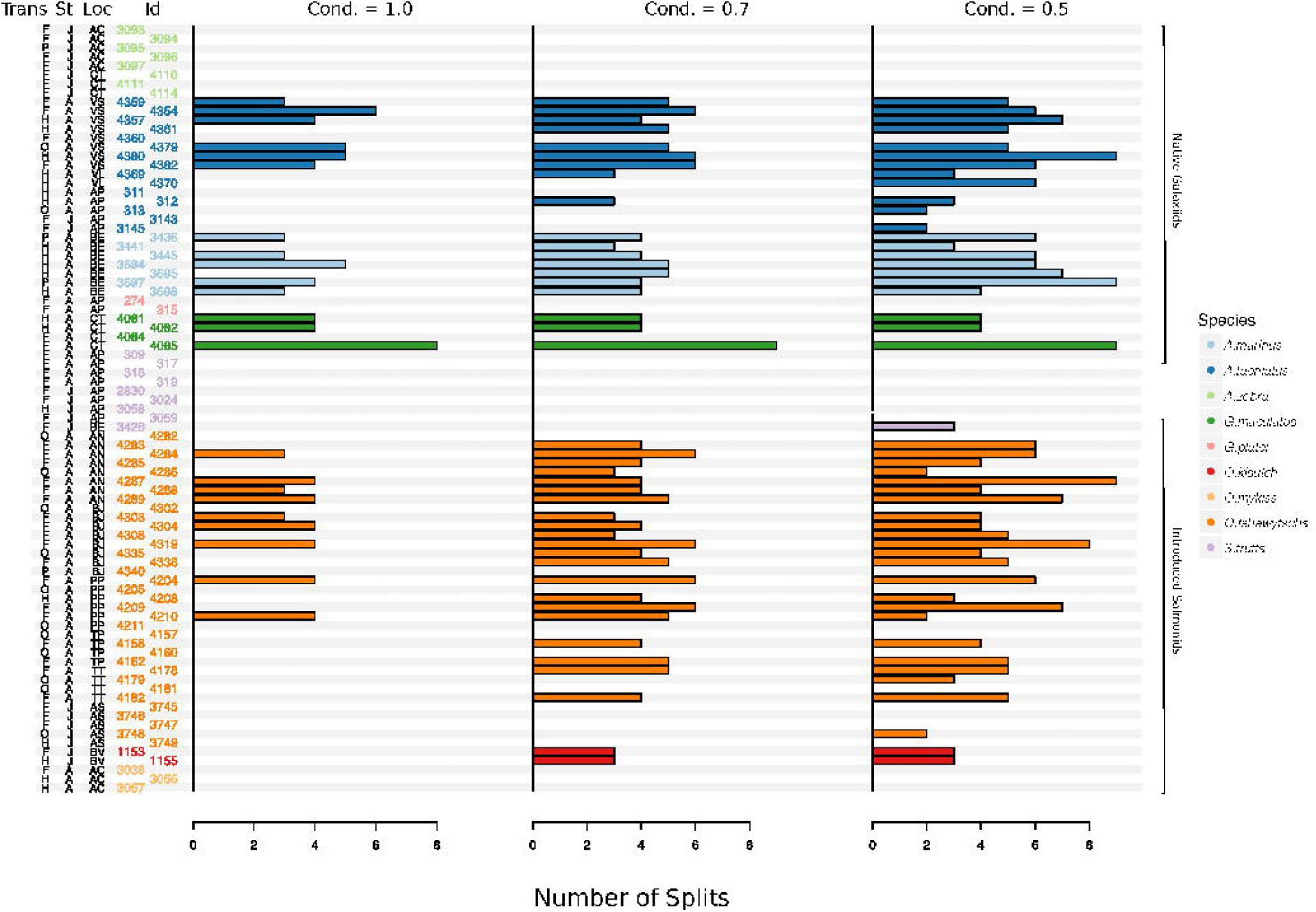
Total number of splits obtained by semisupervised CART on univariate Sr:Ca otolith data for all the species included in the study. Original data for native *A. taeniatus, A. marinus, and G. maculatus* and introduced *O tshawytscha* was divided in more than one homogenous cluster by semisupervised regression trees and led to rejection of the hypothesis of freshwater residency. Details for each individual and split reported in this graph are available in the supplementary material Table S2. “Trans” refers to the quality of the otolith transect, that is: “F” is a full or good quality transect, edge – core – edge, “H” is a half transect, edge – core; “P” is a partial transect, edge – core – extra data without reaching the next otolith edge; “O” is a flagged transect which failed to go through the core and may have some missing data. “St” refers to each fish’s ontogenetic phase at the time of capture, where “J” is for juveniles and “A” for adult specimens; “Loc” indicates the sampling locality where AC: Aysén-Caro, CT: Chiloé-Tocoihue, VS: Valdivia-Santo Domingo, VL: Valdivia-Lingue, AP: Aysén-Palos, BE: Baker-estuary, AN: Aysén-Ñireguao, BJ: Baker-Jaramillo, PP: Petrohué-Patos, TP: Toltén-Peuco, TT: Toltén-Truful, AS: Aysén-Simpson, BV: Baker-Vargas; “ID” is the unique identification of each fish.

### Native galaxiids

Large elemental shifts in otolith profiles indicated a catadromous life-history for most *A. taeniatus* analysed (Table 1 and Fig. 5). Even when confined to strictly freshwater habitats, as in Lake Caro, *A. taeniatus* juveniles showed patterns of habitat mobility (Fig. 5, cond.= 0.5), as contrasted with *A. zebra* or *O. mykiss*, which showed long-term residency in the same environment.

Otolith profiles (Fig. 4 and 5) suggested that *A. marinus* copes with high levels of salinity variation in the Baker River system. Otolith primordia of all specimens of *A. marinus* showed evidence of higher Sr:Ca ratios at early stages of growth, presumably before the fish entered the estuary (site of capture). Taken together, these data suggest that *A. marinus* is catadromous.

Results indicated that *A. zebra* uses a chemically uniform habitat at both collection localities (Fig. 4), although results should be corroborated by future studies because *A. zebra* individuals assayed were juveniles. Nevertheless, specimens from Tocoihue River appear to have been exposed to higher salinity levels than those from Lake Caro (Table S2), suggesting preference for freshwater residency, but capacity for osmoregulation when salinity levels increase.

*G. maculatus* individuals were sampled from the same site as some specimens of *A. zebra* (Tocoihue). However, among the four specimens assayed, three showed exposure to higher salinity than any other galaxiid species analysed (Fig. 3) with evidence of both catadromous and amphidromous transitions from brackish to saltwater (Table 1). The *G. maculatus* specimen in Fig. 4 was caught in the lower reach of Tocoihue River, an area with strong tidal influence. It shows an amphidromous life-cycle with intermediate salinity influence in the primordium followed by a period of residency in higher salinity and subsequent migration back into brackish and freshwater (as also observed by McDowall (1968)). A fourth specimen of *G. maculatus* showed no major Sr:Ca fluctuations across the otolith transect (Table S2), suggesting that this individual did not drift out to the ocean during its larval stage and it was likely a resident of the estuarine area in Tocoihue.

Only one specimen from the low-elevation coastal Palos Lake was assayed for *G. platei*, and results indicate freshwater residency as revealed by uniformly low Sr:Ca ratios across the entire otolith transect (Fig. 3 and 4).

### Introduced salmonids

This study supports established anadromy in *O. tshawytscha*, as previously shown by other authors (Ciancio et al., 2005; Correa & Gross, 2008; Arismendi & Soto, 2012; Araya et al., 2014). Data are consistent with regularly timed changes in Sr:Ca concentration levels that suggest hatching in freshwater, migration to areas with higher salinity concentrations and a return to inland, freshwater areas to spawn (Fig. 4). *O. tshawytscha* specimens collected from the Simpson River do not show as much variation as other *O. tshawytscha* from this study as these fish were all juveniles that had not yet migrated.

The two parr *O. kisutch* analysed revealed one large scale habitat shift between birth and time of capture. Both otolith profiles were characterized by a relatively high Sr:Ca signature around the core which diminished substantially towards the edges (Fig. 4). These specimens where caught during the summer, about 55 km upstream of the Baker River’s mainstream. Thus, observed patterns could plausibly be induced by maternal effects (Kalish, 1990; Volk et al., 2000; Zimmerman & Reeves, 2002). Additionally, there is already some evidence pointing to young-of the-year *O. kisutch* recruits in remote freshwater fjords of southern Chile (Górski et al., 2016). Accordingly, these specimens were interpreted as anadromous by maternal origin. Conversely, *O. mykiss* exhibited a pattern consistent with freshwater residency and minor salinity fluctuations within their habitat over the entire life cycle (Fig. 3 and 5).

Evidence of at least two different life cycle patterns emerged for *S. trutta* specimens caught at three different locations. The Sr:Ca transect of juveniles from Lake Caro and adults from Lake Palos showed a pattern consistent with continuous residency in freshwater (Fig. 3 and 5) whereas *S. trutta* from Baker River showed higher values at the primordium (Fig. 4).

## Discussion

This study quantitatively identified significant transitions across otolith profiles using regression trees on Sr:Ca ratios. Native galaxiids showed considerable variation in habitat shifts when compared across species, with some species exhibiting differences at the population and individual levels, indicating a high degree of plasticity. Of five native galaxiids examined, evidence was found for one catadromous (*A. marinus*) and two facultatively amphidromous or catadromous species (*G. maculatus* and *A. taeniatus*). Nonnative salmonids have established populations with a broad array of migratory life histories, reflective of those found in their native ranges. Patterns consistent with anadromy were present in three (*O. tshawytscha, O. kisutch, S. trutta*) of four species included in this study.

Overall, several species appear to regularly use habitats with different salinity levels. Otolith profiles that showed variation under the most restrictive analytical conditions were those most likely to exhibit large-scale habitat shifts between different environments and salinity levels (*A. taeniatus, A. marinus, G. maculatus, O. tshawytscha*). Otolith profiles that varied under less stringent conditions provided information about subtler shifts within habitat types.

Results suggest a preponderance of euryhaline and facultative diadromous species among native galaxiids and introduced salmonids in Patagonia. The inferences on species migratory status by population reported in Table 1 suggest that some species display a diverse range of life history strategies (facultative diadromy), coinciding with an increasing number of studies reporting flexibility in diadromous patterns for numerous fishes. These studies, which also include some Southern Hemisphere fishes, evidenced the extent of variability of resident/migratory life histories within single species, often shifting from the classical view that tended to categorize species as either exclusively resident or migratory (Hicks et al., 2010; Augspurger et al., 2015; Górski et al., 2015).

Observed otolith microchemistry for introduced salmonids showed that some species have established movement strategies similar to those in their native ranges. The successful establishment of anadromous exotic salmonids in Chile reinforces the hypothesis that the biotic and abiotic conditions required for diadromy to be maintained (Gross et al., 1988) are present in Chilean waters.

Temperate areas of southern Chile may be considered a favourable environment for the development and maintenance of migratory strategy in fishes. While a limited number of fishes around the world are considered to exhibit some form of diadromy (Nelson, 2006), most are confined to areas such as islands (in Iceland, Hawaii, Falklands, Chathams the entire freshwater fish fauna exhibits diadromy) and temperate areas with recent geological origin (McDowall, 1988). Comparative otolith microchemistry analysis suggests that fishes of southern Chile may require a heterogeneous and spatially connected environment to complete their life-cycles. Many species may also have physiological adaptations or plasticity that allow for osmoregulation in a wide range of saline and freshwater environments (euryhaline life history).

### Limitations and further studies

Some otolith results may have been influenced by maternal effects or induced by local temporal variation in water chemistry. Further studies may indicate whether the higher Sr:Ca ratios in primordia observed in some species could be attributable to maternal effects or other causes. For example, although the mechanisms are not completely understood, physiological constraints in early ontogeny could increase the rate of Sr absorption into the calcium carbonate matrix of the otolith (de Pontual et al., 2003). Also, as the Baker corridor is influenced by a large ice field (Campo de Hielo Norte), high amounts of glacier flour (suspended solids) can contribute to increased salinity levels in water that flows into the estuary (Vargas et al., 2011; Marín et al., 2013). These seasonal salinity changes may promote the uptake of Sr into the otolith matrix and confound the assumption of low Sr in freshwater environments (Zimmerman, 2005). Therefore, even though Sr has been traditionally recognized as a very robust marker to discriminate between saline and freshwater environments, several recent studies have indicated that factors as species-specific variation, environmentally-mediated physiological processes and individual variation can influence Sr uptake into the otolith matrix (Sturrock et al., 2015).

This study suggests that considerable variation in migratory life history may exist in Chilean fishes, but its inferential scope is restricted by the limited number of samples which were collected for other research purposes (see Correa & Hendry, 2012; Correa et al., 2012; Alò et al., 2013). Species-specific reference values for Sr:Ca ratios and a more comprehensive sampling will be needed to effectively characterize and quantify the extent of fish movements within Chilean continental waters. Ideally, a variety of different field techniques (natural markers such as stable isotopes, otoliths, statoliths for lampreys, or scales, as well as molecular markers, tagging, trapping and tracking) and laboratory methods (movement physiology, swimming performance, metabolism) (see Dingle, 2014) could be used to characterize daily and seasonal migration patterns. Indeed, experimental methods related to swimming performance and unidirectional movement propensity are now employed in design of fish passage structures to benefit Chilean native fishes (Laborde et al., 2016).

### Conservation issues

This study could help refine the conservation priorities for freshwater fishes in southern Chile. Given high endemism and the likelihood of dependence on diadromous behaviour, potential threats to fishes from fragmentation of river-to-estuary networks are correspondingly high. Hydroelectric power development causes loss of hydrological connectivity and alteration of river flows, disproportionately affecting fishes with migratory life histories.

Comparative otolith microchemistry results underscore the variation in life history strategies that should be accounted for when planning to manipulate water-flow for hydroelectric developments. Diadromous species depend on the habitat diversity and complexity created by unobstructed watersheds and are locally extirpated when barriers preclude movement to essential habitat.

Additionally, anthropogenic barriers and alterations to water flow (e.g., hydropeaking) may also negatively affect landlocked populations because such structures disrupt successful reproduction, recruitment and habitat quality (Alò & Turner, 2005; Fullerton et al., 2010; Garcia et al., 2011). Current hydropower capacity in Chile, now ∼6.000 megawatts of energy connected to the central grid, is expected to nearly double to ∼ 11.000MW by the year 2020 in the southern sector of the country (Santana et al., 2014). Development is slated in basins that harbour the majority of native fish species diversity.

Ongoing spread of exotic species threatens native species through negative interactions including predation, competition, behavioural inhibition and homogenization (Correa & Gross, 2008; Penaluna et al., 2009; Correa & Hendry, 2012; Correa et al., 2012; Habit et al., 2012; Arismendi et al., 2014; Vargas et al., 2015). In particular, establishment of migration runs of *O. tshawytscha* and *O. kisutch* could trigger additional threats such as the shift of significant amounts of marine-derived nutrients to previously oligotrophic environments (Helfield & Naiman, 2001; Arismendi & Soto, 2012) and increased competition for limited resources with the native diadromous species. Ironically, non-native salmon and trout are also likely to be negatively affected by future hydroelectric dams. Additional hydropower development will almost certainly impact a flourishing tourism industry supported by salmonid recreational fisheries (Arismendi & Nahuelhual, 2007; Vigliano et al., 2007).

The evolutionary processes that allowed dispersal and colonization of Patagonian fishes are influenced by the region’s unique geography, climate, and geological processes. To ensure proper conservation of native freshwater, diadromous, and commercially relevant sport fisheries, managers will have to carefully designate and protect critical habitats, and in many cases mitigate obstruction of river flows imposed by dams with appropriate fish passage structures (Wilkes et al., 2017). Long-term monitoring should also be a priority to understand the broad impacts of hydropower development on aquatic biodiversity.

## Supporting Information

All elemental data will be provided in a data dryad file (http://datadryad.org/) upon publication of this manuscript. The following additional tables and figures are provided as supplement to this manuscript and are available for review:

Table S1: A list of Continental Native Fishes of Chile, including life histories and/or habitat.

Table S2: Description of the samples analysed and CART Splitting means and standard deviations.

S3: Pictures of each otolith associated with a graph reporting Sr:Ca transect data.

## Acknowledgements

We are thankful to A.P. Bravo, I.Y. Quinteros, M. Soto-Gamboa, and L. Caputo for assistance in the field, and to M.L. Guillemin for providing field equipment. S. Platania and M. Brandenburg assisted with laboratory procedures and logistics. The Museum of Southwestern Biology at UNM provided curatorial assistance. We thank P. Marquet for revising an earlier version of this manuscript, A. Castillo for help with GIS mapping, E. Habit for a useful discussion.

